# Conversion between 100-million-year-old duplicated genes contributes to rice subspecies divergence

**DOI:** 10.1101/2020.12.22.424042

**Authors:** Chendan Wei, Zhenyi Wang, Jianyu Wang, Jia Teng, Shaoqi Shen, Qimeng Xiao, Shoutong Bao, Yishan Feng, Yan Zhang, Yuxian Li, Sangrong Sun, Yuanshuai Yue, Chunyang Wu, Yanli Wang, Tianning Zhou, Wenbo Xu, Jigao Yu, Li Wang, Jinpeng Wang

## Abstract

Extensive sequence similarity between duplicated gene pairs produced by paleo-polyploidization may result from illegitimate recombination between homologous chromosomes. The genomes of Asian cultivated rice Xian/*indica* (XI) and Geng/*japonica* (GJ) have recently been updated, providing new opportunities for investigating on-going gene conversion events. Using comparative genomics and phylogenetic analyses, we evaluated gene conversion rates between duplicated genes produced by polyploidization 100 million years ago (mya) in GJ and XI. At least 5.19%–5.77% of genes duplicated across three genomes were affected by whole-gene conversion after the divergence of GJ and XI at ~0.4 mya, with more (7.77%–9.53%) showing conversion of only gene portions. Independently converted duplicates surviving in genomes of different subspecies often used the same donor genes. On-going gene conversion frequency was higher near chromosome termini, with a single pair of homoeologous chromosomes 11 and 12 in each genome most affected. Notably, on-going gene conversion has maintained similarity between very ancient duplicates, provided opportunities for further gene conversion, and accelerated rice divergence. Chromosome rearrangement after polyploidization may result in gene loss, providing a basis for on-going gene conversion, and may have contributed directly to restricted recombination/conversion between homoeologous regions. Gene conversion affected biological functions associated with multiple genes, such as catalytic activity, implying opportunities for interaction among members of large gene families, such as NBS-LRR disease-resistance genes, resulting in gene conversion. Duplicated genes in rice subspecies generated by grass polyploidization ~100 mya remain affected by gene conversion at high frequency, with important implications for the divergence of rice subspecies.

**One-sentence summary:** On-going gene conversion between duplicated genes produced by 100 mya polyploidization contributes to rice subspecies divergence, often involving the same donor genes at chromosome termini.

## Introduction

Rice is the largest food crop in the world. There are two distinct types of domesticated rice, Asian rice (*Oryza sativa*) and African rice (*Oryza glaberrima*), each with unique histories of domestication (Sweeney and McCouch, 2007). Asian rice is planted worldwide, feeding half of the world’s population as staple food and providing more than 20% of the energy for human survival (Kim et al., 2008, Stein et al., 2018, Wang et al., 2018b). Xian/*Indica* (XI) and Geng/*Japonica* (GJ) are the two major subspecies of rice, which diverged ~0.4 million years ago (mya). The first whole-genome draft sequence of GJ cultivar ‘Nipponbare’, which is representative of the subspecies, was obtained in 2002 (Goff et al., 2002), and genome sequencing and annotation have been continuously improved (Tanaka et al., 2008). The whole-genome sequence of XI (93-11) has also been deciphered (Yu et al., 2002), with high-quality genome sequences of representative varieties Zhenshan 97 (XI-ZS97) and Minghui 63 (XI-MH63) made available (Zhang et al., 2016). These two main varieties of XI are the parents of an excellent Chinese hybrid. XI accounts for more than 70% of global rice production and possesses much higher genetic diversity than GJ (Huang et al., 2010), as highlighted by recent analysis of 3,010 diverse Asian cultivated rice genomes and 1,275 rice varieties (Li et al., 2020, Wang et al., 2018b).

Recursive polyploidization or whole-genome duplication (WGD) is the doubling of an entire set of chromosomes in cells and is prevalent throughout the plant and animal kingdoms (Frawley and Orr-Weaver, 2015). The impact of polyploidization on plant functional evolution is extremely profound, facilitating rapid expansion and divergence of species (Jiao et al., 2011, Puchta et al., 1996, Barker et al., 2016, Wu et al., 2020). A large number of duplicated genes generated by polyploidization are distributed on the homologous chromosomes of extant species, which leads to genome instability. Homoeologous recombination may result in loss of large segments of DNA (Paterson et al., 2004, Zhuang et al., 2019), de novo functionalization of genes, subfunctionalization (Taylor and Raes, 2004), or rearrangement of genomic DNA (Wang et al., 2005, Murat et al., 2017, Wang et al., 2018a, Wang et al., 2017), providing material for plant evolution. At least five WGD events occurred during the formation of modern cultivated rice. The two oldest are a WGD event (called ζ) shared by seed plants (divergence ~310 mya) and a WGD event (called ε) that occurred prior to the appearance of the most recent common ancestor of all extant angiosperms (~235 mya) (Jiao et al., 2011). Two relatively recent WGD events occurred after the formation of monocotyledons: one (τ) shared by most monocotyledons at ~130 mya, and another (σ) shared by Poales at ~115-120 mya (Tang et al., 2010, Paterson et al., 2004, Ming et al., 2015). The most recent WGD event (ρ) was originally thought to have occurred before the divergence of major grasses (~70 mya) (Paterson et al., 2004, Wang et al., 2005); however, the latest fossil evidence advances this ρ event to ~100 mya (Wang et al., 2015).

Homologous recombination provides a major source of genetic innovation (Kurosawa and Ohta, 2011). In plants, meiotic and mitotic recombination result in reciprocal or symmetric exchange of DNA sequence information between homologous chromosomes (Gardiner et al., 2019). In addition to normal genetic recombination, highly similar sequences undergo frequent recombination between homologous chromosomes, which is called illegitimate recombination (Wang et al., 2009). One result of this recombination is gene conversion, where one gene (or DNA fragment) replaces another gene (or DNA fragment) on a homologous chromosome or chromosomal region. Gene conversion between duplicated genes produced by polyploidization has been identified in the genomes of *Poaceae*, *Arachis hypogaea*, *Gossypium*, *Brassica campestris*, and *Brassica oleracea* (Wang et al., 2009, Wang et al., 2011, Paterson et al., 2012, Zhuang et al., 2019, Yu et al., 2013, Liu et al., 2020). Gene conversion is frequent and on-going between homologous chromosomes, such as homologous chromosomes 11 and 12 produced from the duplication common to grasses (ρ event) in the modern rice genome (Wang et al., 2009, Kurosawa and Ohta, 2011, Wang and Paterson, 2011, Wang et al., 2019).

Recombination is a mutagenic factor, and mutations lay the foundation for natural selection. The main role of gene conversion is to maintain the homology or similarity of duplicated sequences. Comparison between rice and sorghum clearly suggests that gene conversion promotes gene divergence (Wang et al., 2009). Recombination accelerates mutation, with gene conversion playing an important role (Guo et al., 2013). Gene conversion of functional sequences and new mutations produced by related homologous recombination may affect gene function. Sequences encoding functional domains are converted more frequently than those encoding non-functional domains (Wang et al., 2007). Gene conversion and DNA duplication may facilitate functional innovation through gene extension and mutations in structural domains of disease-resistance genes (Ratnaparkhe et al., 2011). Gene conversion between chromosomes 11 and 12 of rice has been accompanied by subfunctionalization or purifying selection of genes related to spikelet abortion (Zhang et al., 2011), lipid transfer genes (Jang et al., 2008, Wang et al., 2012), two recessive yellowing control genes (Mao et al., 2011), genes encoding cyclic C2-type proteins (Jung et al., 2012), and the zinc-inducible promoter family (Ricachenevsky et al., 2011).

Our knowledge of gene conversion between paralogous genes in the two rice subspecies (Wang et al., 2007) is based on outdated genomic data (ver. 4), and the imperfections in genome sequencing assembly and annotation particularly may have implications for gene conversion analysis. Here, we used the latest genomic data and recent approaches for resolving genomic homology (Wang et al., 2018a) to identify paralogous genes generated by WGD event (ρ) in three rice genomes representing the two major subspecies. We then combined this with comparative and phylogenetic genomics to establish an improved method for inferring gene conversion. We evaluated the ratio, level, and pattern of gene conversion in three rice genomes and explored the effects of this process on genome evolutionary rate, gene function innovation, chromosome structure, and genome stability.

## Results

### Intra/intergenomic homologous genes

We performed genomic colinearity and structure analysis and identified duplicated genes generated by the WGD event common to grasses in the GJ, XI-MH63, and XI-ZS97 genomes. For blocks containing more than four colinear genes, there were more duplicated genes in GJ (3,314 pairs) than in XI-MH63 or XI-ZS97 (2,629 pairs and 2,889 pairs, respectively). We identified 46, 18, and 10 homologous blocks with more than 10, 20, and 50 colinear gene pairs in GJ, respectively. XI genomes had much shorter duplicated blocks, with fewer than 10 blocks possessing more than 50 colinear gene pairs (**Supplemental Table 1**). We also used a bidirectional best BLAST homology search to identify homologous gene pairs residing in paralogous regions because some pairs might have been removed from the colinearity analysis. Finally, 3,256, 2,502, and 2,816 homologous gene pairs were identified in GJ, XI-MH63, and XI-ZS97, respectively (**Figure 1**). Compared with GJ, the XI varieties had fewer homologous genes because XI has experienced more chromosomal rearrangement events.

**Figure 1.**
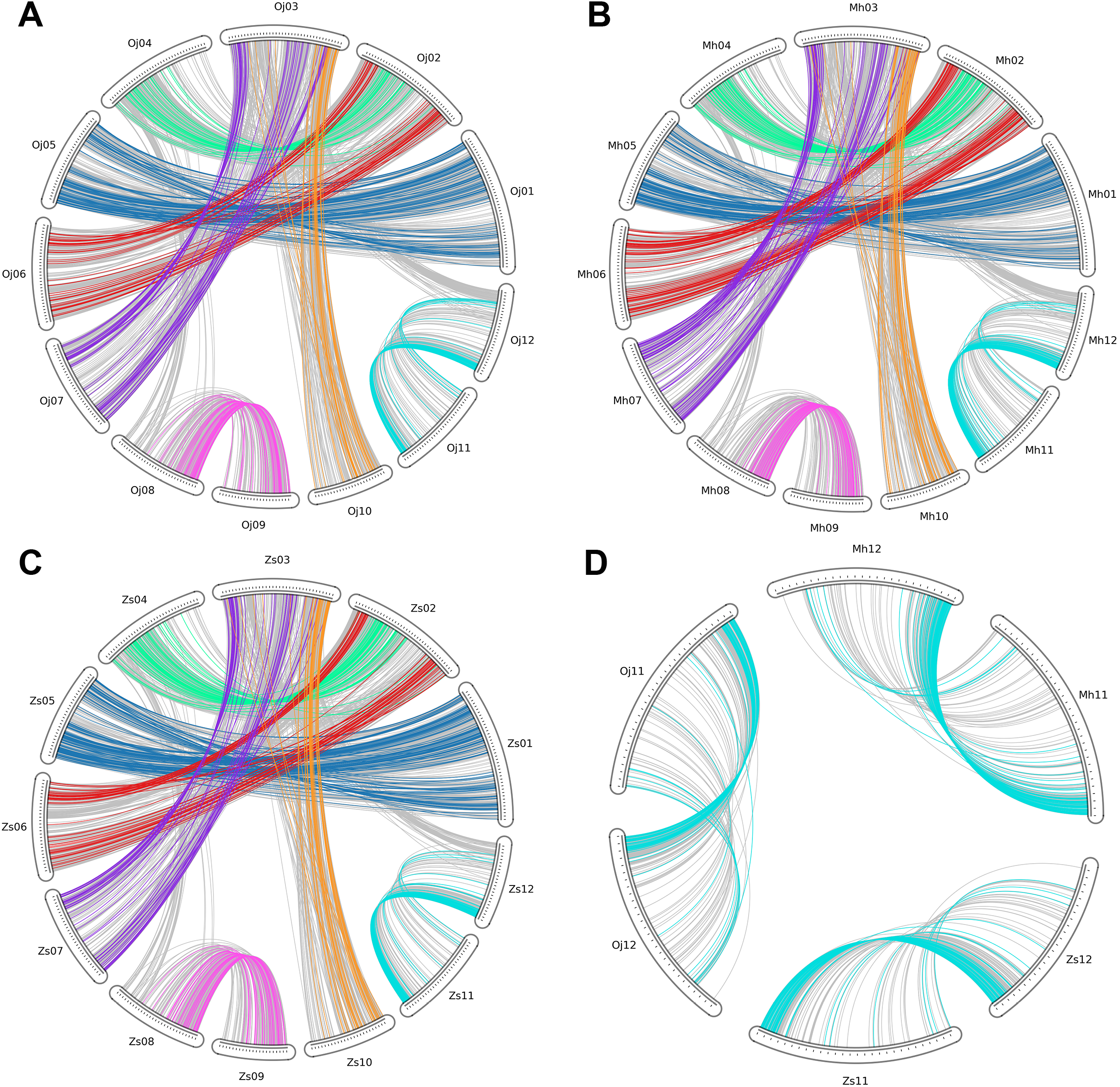
Genome duplications and conversion patterns in three rice subspecies genomes. Lines show duplicated gene pairs between chromosomes in three genomes. Colored lines indicate gene-conversion pairs; grey lines indicate non-gene-conversion pairs. (A) Gene duplication and gene conversion in GJ. (B) Gene duplication and gene conversion in XI-MH63. (C) Gene duplication and gene conversion in XI-ZS97. (D) Gene duplication and gene conversion on chromosomes 11 and 12 of GJ, XI-MH63, and XI-ZS97.

We used colinearity and structure analysis of intergenomic homologous genes to infer orthologous genes generated by the recent species divergence (**Supplemental Table 1**). Colinearity analysis identified 19,089 orthologous gene pairs in 103 blocks between the GJ and XI-MH63 genomes. Between GJ and XI-ZS97, there were 18,498 orthologous gene pairs in 119 blocks. The two varieties of XI, XI-MH63 and XI-ZS97, showed better colinearity, with 25,262 orthologous gene pairs between them in 146 blocks. We again performed a bidirectional best BLAST homology search among the three genomes to identify additional orthologous genes. There were 23,719 orthologous gene pairs between GJ and XI-MH63, and 23,056 orthologous gene pairs between GJ and XI-ZS97. Since XI-MH63 and XI-ZS97 are more closely related, we identified 35,049 orthologous gene pairs between the genomes of these varieties (**Supplemental Table 2**).

### Homologous gene quartets

To detect possible gene conversion between homologous genes produced by WGD, we used homology and colinearity information to identify homologous gene combinations for WGD and species divergence, which we defined ‘homologous gene quartets.’ Assuming that the genomes of two species (‘O’ and ‘S’) retain a pair of duplicated chromosomal generated in a common ancestor through WGD, the paralogous genes O1 and O2 in species O and the respective orthologous genes S1 and S2 in species S constitute a homologous gene quartet (**Figure 2A**). Sequence similarity between orthologous gene pairs is more similar than that between paralogous gene pairs if there is no gene conversion (or nonreciprocal recombination) between the duplicated gene pairs after species divergence (**Figure 2B**). However, if gene conversion occurs between duplicated genes, we might find that the gene tree topology has a different structure than expected (**Figure 2C-E**). Changes in the topological structure of the gene tree can be determined from the similarity of homologous sequences in homologous gene quartets. As the gene sequence may be converted in whole or in part, we used different methods to infer whole-gene conversion (WCV) and partial-gene conversion (PCV) (see Materials and Methods for details).

**Figure 2.**
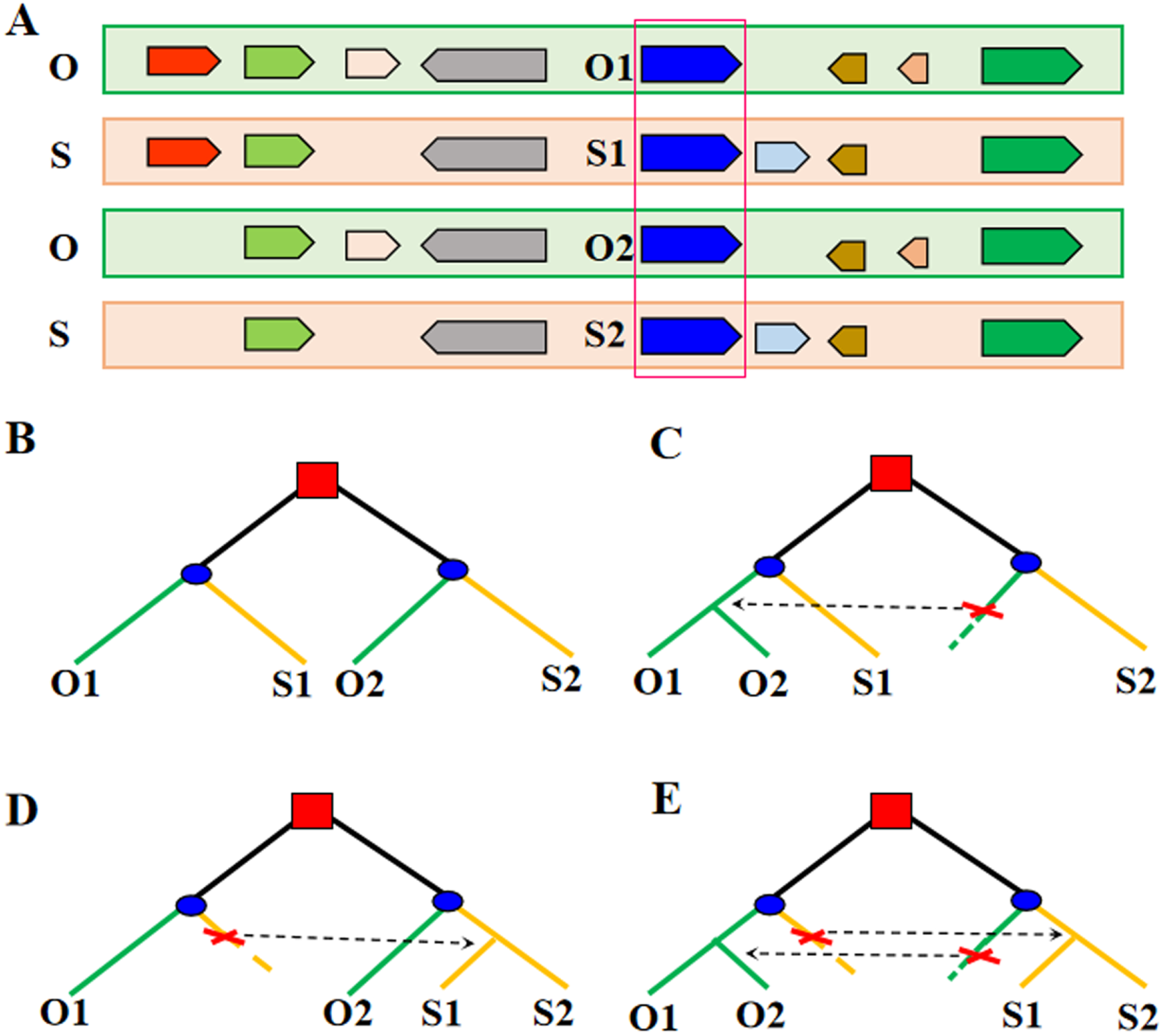
Gene conversion events were inferred by construction of homologous gene quartets and changes in phylogenetic tree topology. (A) Colinear chromosomal segments from two genomes (O and S), represented by rectangles of different colors. Arrows show genes, and homologous genes are indicated by the same color. Homologous gene quartets are formed by paralogous genes O1 and O2 in one genome and their respective orthologs S1 and S2 in the other genome. (B-E) Squares symbolize a WGD event in the common ancestral genome; circles symbolize species divergence. (B) The expected phylogenetic relationship of the homologous genes if no conversion occurs. (C) O2 (an acceptor) is converted by O1 (a donor). (D) S1 is converted by S2. (e) Both of the above conversions occur.

Based on colinearity information of intragenomic and intergenomic homologous genes, we identified 2,788 quartets between GJ and XI-MH63, and 2,879 quartets between GJ and XI-ZS97. Although XI-MH63 and XI-ZS97 are varieties of the same subspecies, relatively few quartets (2,566) were identified between them, probably due mainly to differences in gene loss after the three genomes diverged. By comparing the three genomes, we inferred a possible ancestral gene content before divergence of 19,104. Rates of gene loss or translocation were 6.13%, 13.31%, and 7.89% in GJ, XI-MH63, and XI-ZS97, respectively. Finally, we identified 3,332, 3,322, and 3,254 homologous genes in GJ, XI-MH63, and XI-ZS97, respectively. These homologous genes were mainly conserved in 82, 85, and 93 blocks, and they were unevenly distributed across the 12 chromosomes in the three genomes (**Figure 1**).

### Gene conversion and occurrence patterns

We removed highly divergent sequences to reduce the possibility of inferring gene conversion events from unreliable sequences (see Materials and Methods for details). After this, 2,788 gene quartets were identified between GJ and XI-MH63, 2,879 quartets between GJ and XI-ZS97, and 2,566 quartets between XI-ZS97 and XI-MH63 (**Supplemental Table 2**). We used two methods to infer gene tree topology, one based on synonymous nucleotide substitution rate (*Ks*) as a similarity measure and the other based on amino acid identity ratio, which we called whole-gene conversion type I and type II (WCV-I and WCV-II), respectively. We used a combination of dynamic planning and phylogenetic analysis to infer possible partial-gene conversion (PCV) events (**Supplemental Table 3**). Since paralogous gene pairs may be identified in different quartets, we merged the paralogous gene pairs affected by gene conversion in each genome. This gave us the gene conversion events of each genome after the divergence of rice.

In GJ, 398 pairs (~12%) of paralogs had been converted. Of these, 179 pairs (5.37%) had undergone WCV: 11 pairs were inferred byWCV-I and 168 pairs were inferred by WCV-II. Another 259 pairs (7.77%) were PCVs, which occurred at a remarkably higher rate than WCV. In XI-MH63, 466 pairs (~14%) of paralogs had been converted, of which 182 pairs (5.48%) were WCVs: 8 pairs were inferred by WCV-I and 174 pairs were inferred by WCV-II. Another 312 pairs (9.39%) were PCVs, which was significantly higher than WCVs. In XI-ZS97, 468 pairs (~14%) of paralogs had been converted: 185 pairs (5.69%) were WCVs, comprising 8 pairs inferred by WCV-I and 177 pairs inferred by WCV-II. Another 310 pairs (9.53%) were PCVs, which was also more than the number of WCVs (**Table 1**). For example, we detected gene conversion between the paralogous genes *Zs11g0407.01* and *Zs12g0396.01*, and one gene fragment from 335 to 462 bp was converted through one-way genetic information transmission (or rearrangement) (**Figure 3A**). We discovered that the gene conversion rate in XI was significantly higher than that in GJ (**Figure 3B**). By analyzing topological changes in the gene trees reconstructed using homologous genes, we further determined that gene conversion occurred between *Mh11g0214.01* and *Mh12g0189.01* (**Figure 3C; Supplemental Text**).

**Table 1.**
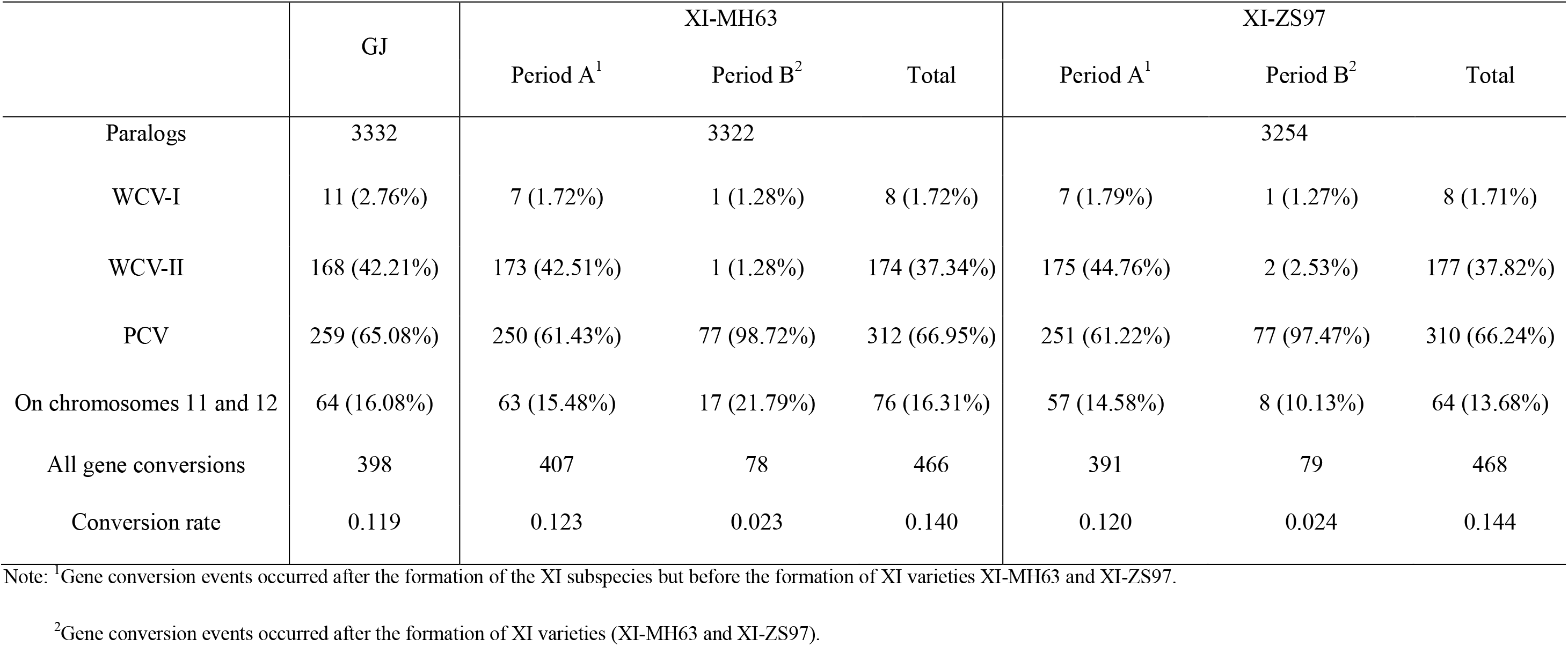
Converted paralogs in GJ and XI genomes (XI-MH63 and XI-ZS97).

**Figure 3.**
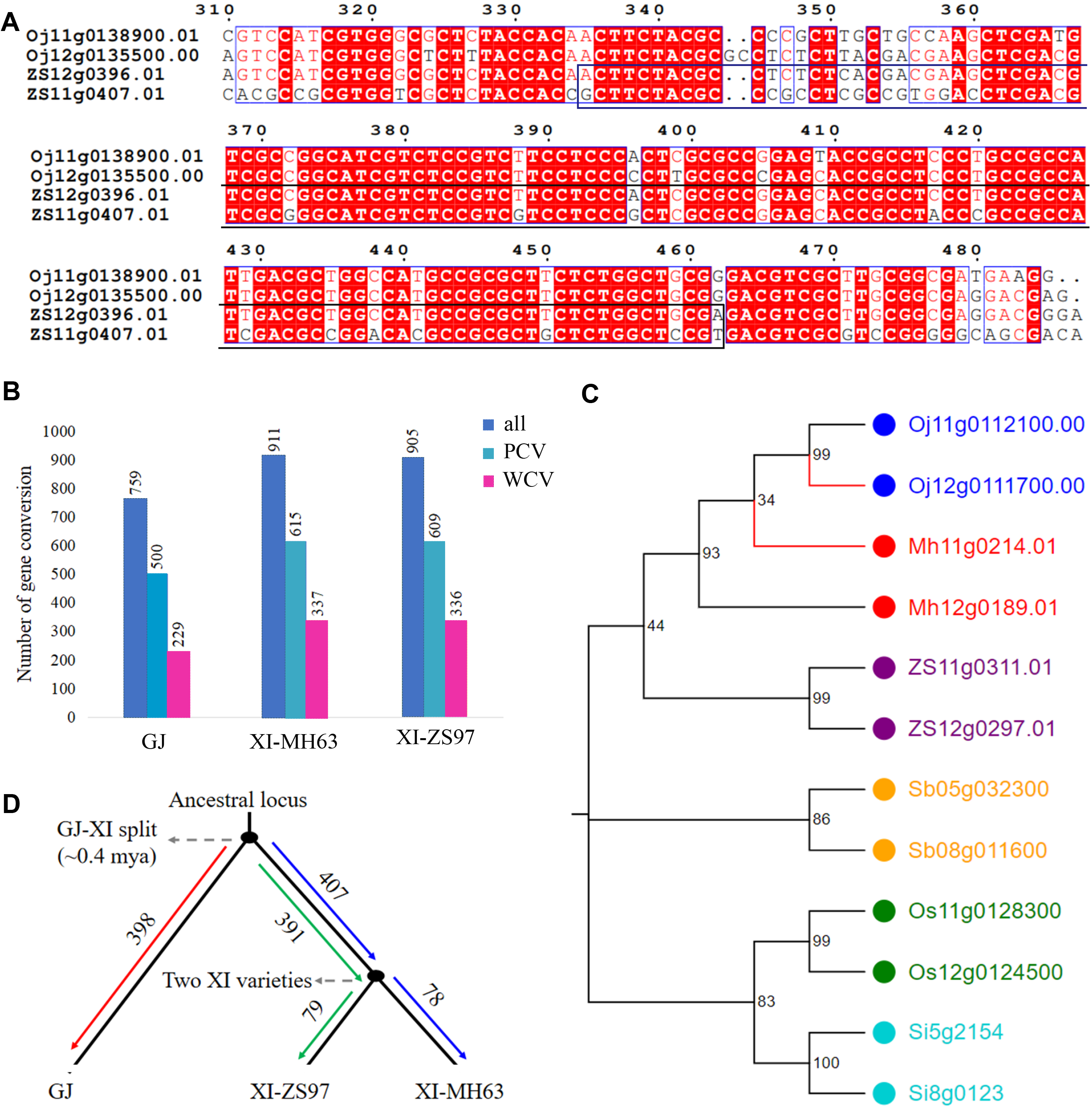
Evolution of gene conversion. (A) Sequence alignment for a homologous gene quartet. The nucleotide sequence from 335 to 462 bp of *Zs12g0396.01* and *Zs11g0407.01* has undergone gene conversion, with *Zs11g0407.01* as the donor. (B) The number of WCV and PCV events occurring in the three genomes. (C) Evolutionary tree of genes in which gene conversion has occurred. the numbers at nodes represent boostrap value. Gene conversion has occurred in *Mh11g0214.01* and *Oj12g0111700.00*. (D) Gene conversion in species divergence events.

### High-frequency on-going gene conversion

By comparing the similarity of homologous gene quartets between different genomes, we inferred gene conversion events in the three genomes during different evolutionary periods. Duplicated gene pairs produced ~100 mya were still being affected by gene conversion. In GJ, we identified 398 pairs of paralogous genes that might have undergone gene conversion after the divergence of rice (**Figure 3D**). The amino acid identity of four (1.01%) pairs of paralogous genes was > 99%, with *Ks* < 0.01 (**Supplemental Figures 1 and 2**). A relatively large number of duplicated genes were affected by gene conversion in the two XI varieties, XI-MH63 and XI-ZS97. In XI-MH63, we found 466 pairs of paralogous genes that might have undergone gene conversion (**Figure 3D**) after the divergence of rice subspecies; six (1.29%) of these pairs of paralogous genes had > 99% amino acid identity between them and *Ks* < 0.01 (**Supplemental Figures 1 and 2**). Similarly, we identified 471 pairs of paralogous genes in XI-ZS97 that might have undergone gene conversion after GJ diverged from XI (**Figure 3D**), and six (1.27%) of these pairs of paralogous genes had > 99% amino acid identity between them and *Ks* < 0.01 (**Supplemental Figures 1 and 2**). We identified small synonymous and nonsynonymous nucleotide substitutions and high sequence identity between duplicated gene pairs in which gene conversion had occurred, suggesting that gene conversion may have occurred over a very short time.

Another striking indication was that 407 and 391 pairs of paralogous genes were affected by gene conversion before the formation of XI-MH63 and XI-ZS97, respectively; 78 and 79 pairs of paralogous genes were converted after formation of the two varieties, accounting for 16.7% and 16.6% of the total gene conversion, respectively (**Figure 3D**). Duplicated genes in XI-MH63 and XI-ZS97 sharing a homologous region showed nearly 99% amino acid identity and 0.99 nonsynonymous nucleotide substitution rate (*Ks*) (**Supplemental Figures 1 and 2**). These data suggest that gene conversion between paralogous gene pairs is on-going and occurs at high frequencies in rice subspecies.

### A donor is usually a donor

Gene conversion involves a donor locus and an acceptor locus. Donors and acceptors can be identified by comparing topological changes in the phylogenetic trees of homologous gene quartets since the paralog of the donor should be more similar than its ortholog. Donors have at least 30% more converted sites than acceptors. We found that 765, 934, and 930 duplicated genes had been converted in GJ, XI-MH63, and XI-ZS97, respectively, with 196, 215, and 200 of these representing donors. A total of 1,520 duplicated genes had been converted in the three genomes, with 1,378 (90.66%) of these converted in two or three genomes. Interestingly, 113 (88.98%) genes had preferred donors in at least two genomes, and 85 (66.93%) genes had the same donor in the three genomes (**Supplemental Table 4**). This suggested that the duplicated gene that had undergone gene conversion was usually present as a donor locus in each different genome (**Figure 4A**). For example, in the region of ~1.0 Mb near the telomere on chromosomes 11 and 12, gene conversion had occurred in 13 duplicated genes. Twelve duplicated genes had undergone gene conversion in at least two genomes. Ten duplicated genes were present as donors, and seven duplicated genes acting as donors in different genomes (**Figure 4B**).

**Figure 4.**
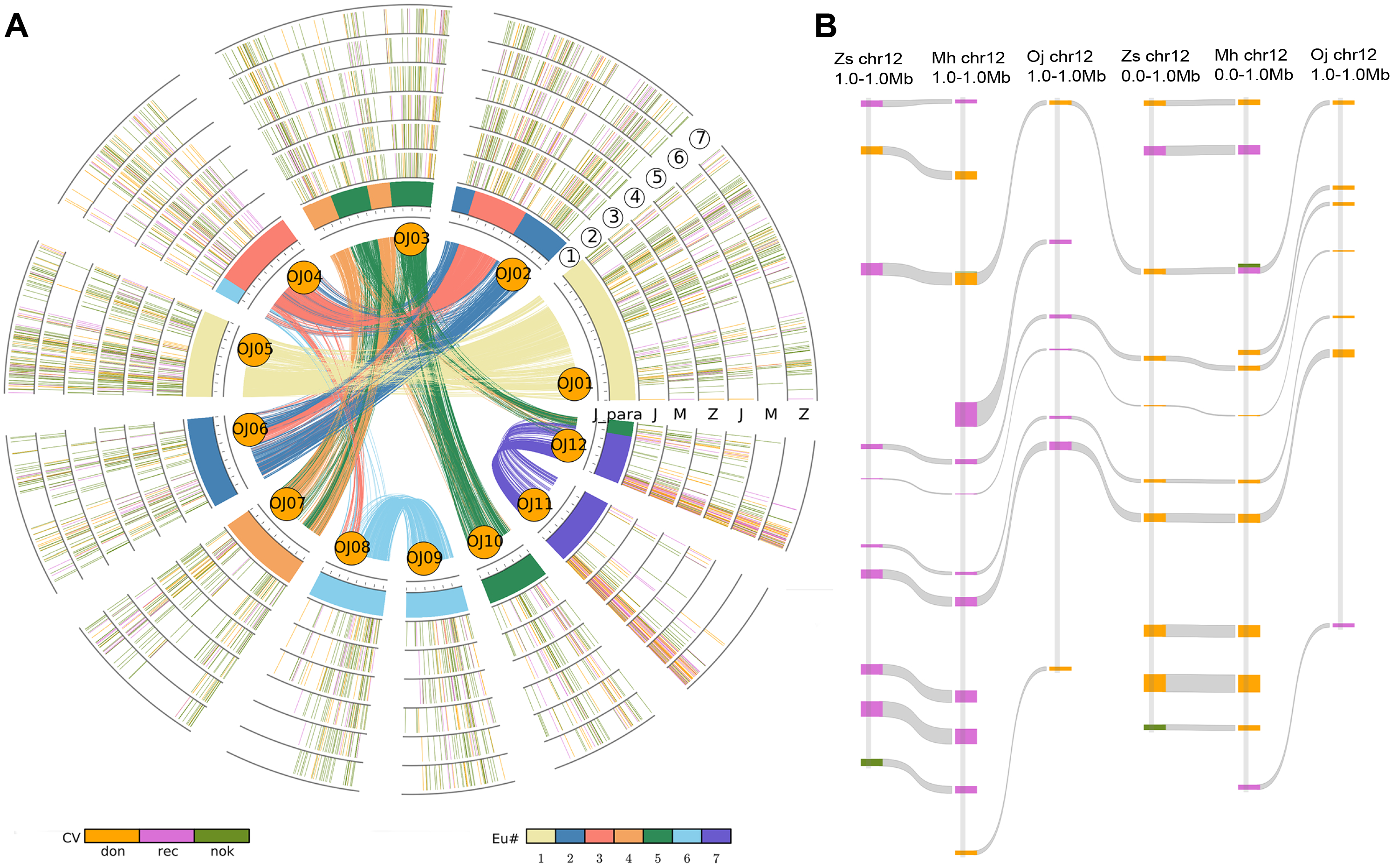
Distribution of donors and receptors in the genome where gene conversion occurs. (A) Homologous distribution of donors and acceptors on chromosomes undergoing gene conversion. Curved lines within the inner circle are formed by 12 chromosomes color coded to the seven ancestral chromosomes before the WGD event common to grasses (ECH) (Wang et al., 2015). Intra-loop curves show duplicated gene pairs in GJ. The inner three circles show the relationships of orthologous gene distribution between the three genomes in which gene conversion has occurred. The outer three circles show the distribution between the three genomes undergoing gene conversion, and the inner three circles show paralogous homologs. Different colors indicate donor (orange) or acceptor (pink) loci, as well as some uncertain loci (green). (B) Local gene conversion and the distribution of donor and acceptor loci. Pink swatches represent donor loci, orange swatches represent acceptor loci, and green swatches represent those loci where donor or acceptor status is uncertain. And Zs means XI-ZS97; Mh means XI-MH63; Oj means GJ.

### Gene conversion and uneven distribution

Gene replacement and conversion were unevenly distributed across the different paralogous homologous chromosomal regions, and all three genomes were most affected by gene replacement and conversion between duplicated genes on chromosomes 11 and 12. The gene conversion rate was 18.88%, 21.78%, and 18.71% on chromosomes 11 and 12 of GJ, XI-MH63, and XI-ZS97, respectively (**Supplemental Table 5**). In GJ, XI-MH63, and XI-ZS97, gene conversions were clustered in the 2 Mb region at the termini of chromosomes 11 and 12, and the gene conversion rate was 74.60%, 67.11%, and 73.02%, respectively. This suggests that gene conversion usually occurs at the termini of chromosomes. (**Figure 1D**).

The physical location of genes on chromosomes may influence the chance of gene conversion. Gene conversion is usually found at the termini of chromosomes, where gene density is high (**Figure 1; Table 2**). In GJ, 692 paralogs were located in the 2 Mb at the termini of chromosomes and about 17.20% of the paralogs were converted. This was higher than the gene conversion rate for the whole genome (12.09%). In XI-MH63, we found 584 paralogs in the 2 Mb at the termini of chromosomes, and approximately 25.34% showed gene conversion, which was also higher than the gene conversion rate for the whole genome (18.57%). In XI-ZS97, there were 675 paralogs located in the 2 Mb close to the termini of chromosomes, of which about 20.59% had undergone gene conversion, which was higher than the gene conversion rate for the whole genome (16.62%). We found that the physical location of genes on chromosomes may correlate with the chance of gene conversion, with genes near the chromosomal termini more frequently affected by gene conversion.

**Table 2.** Relationship between gene physical location and gene conversion

| Distance to telomere | <2 Mbp | 2-4 Mbp | 4-6 Mbp | 6-8 Mbp | 8-10 Mbp | >10 Mbp | Total |
| --- | --- | --- | --- | --- | --- | --- | --- |
| GJ |  |  |  |  |  |  |  |
| All converted | 119 (17.20%) | 60 (13.02%) | 71 (15.78%) | 25 (8.42%) | 12 (8.45%) | 478 (11.16%) | 765 (12.09%) |
| Paralogous genes | 692 | 461 | 450 | 297 | 142 | 4283 | 6326 |
| XI-MH63 |  |  |  |  |  |  |  |
| All converted | 148 (25.34%) | 74 (20.00%) | 68 (23.78%) | 35 (18.52%) | 8 (7.92%) | 576 (17.11%) | 909 (18.57%) |
| Paralogous genes | 584 | 370 | 286 | 189 | 101 | 3366 | 4896 |
| XI-ZS97 |  |  |  |  |  |  |  |
| All converted | 139 (20.59%) | 76 (16.03%) | 70 (18.57%) | 37 (14.98%) | 12 (9.16%) | 578 (16.14%) | 912 (16.62%) |
| Paralogous genes | 675 | 474 | 377 | 247 | 131 | 3581 | 5486 |

### Effect of chromosome rearrangement on gene conversion

Chromosome rearrangement is a random process, and block number in the genome can reflect the degree of chromosome rearrangement after polyploidization. Block number and gene conversion rate showed a positive correlation (**Supplemental Table 6**) in XI-MH63 (R^2^ = 0.22, P-value = 0.12), XI-ZS97, and GJ. However, there was no significant positive correlation in the three genomes (**Figure 5A**). If four special homologous chromosomes (homologous chromosome pairs 1-5 and homologous chromosome pairs 11-12) were removed, there was a significant positive correlation between block number and gene conversion rate in XI-MH63 (R^2^ = 0.85, P-value < 0.01). There was also a significant positive correlation between block number of the chromosomes and gene conversion rate in XI-ZS97 (R^2^ = 0.75, P-value < 0.01) and GJ (R^2^ = 0.74, P-value < 0.01) (**Figure 5B**).

**Figure 5.**
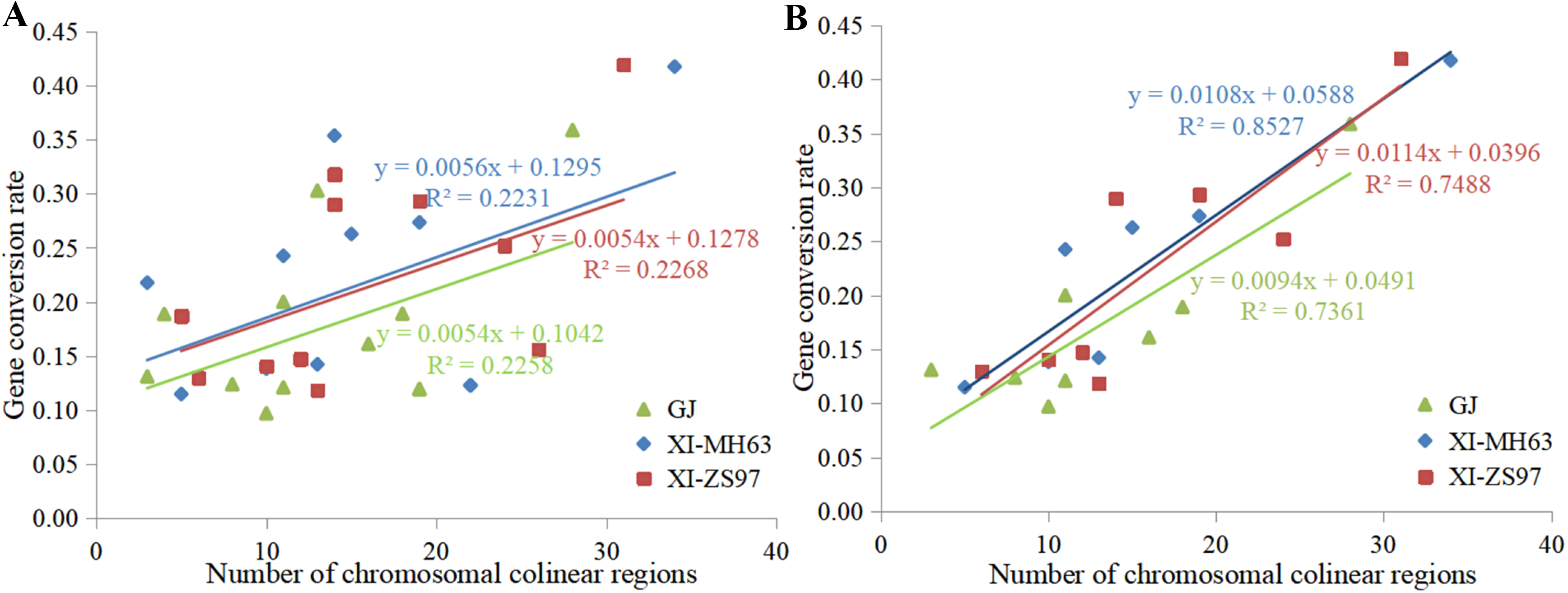
Relationship between block number and gene conversion rate on each chromosomes. (A) Relationship between block number on 12 chromosomes and gene conversion rate on the corresponding chromosomes of GJ, XI-MH63, and XI-ZS97. (B) Relationship between block number on 8 chromosomes and gene conversion rate on the corresponding chromosomes after removing the four special chromosomes (homologous chromosome pair 1-5 and homologous chromosomes pair 11-12).

Correlation does not imply a direct factor leading to gene conversion. For this reason, we further analyzed the relationship between block length and gene conversion rate on each chromosome (**Supplemental Table 7**). We found that longer blocks had a higher gene conversion rate (**Supplemental Figure 3**). The average gene conversion rate for a total of 14 blocks with more than 100 paralogous gene pairs was 14.12% (349 pairs). The block with fewer than 20 paralogous gene pairs was block 219, with a gene conversion rate of 11.77% (178 pairs). These results indicate that the direct result of chromosome rearrangement is the loss of duplicated genes, which may increase the chances of gene conversion. However, chromosome rearrangement may also reduce recombination between chromosomes and inhibit gene conversion.

### Gene conversion and evolution

Gene conversion homogenizes paralogous gene sequences. This makes the affected homologous genes appear younger than expected, based on sequence divergence with one another. The synonymous substitution rate (Pn) and nonsynonymous substitution rate (Ps) between paralogs undergoing gene conversion were smaller than those of paralogs not affected by gene conversion **(Table 3)**. In GJ, the average Pn=0.20 and Ps=0.46 for converted genes were significantly smaller than the average Pn=0.25 and Ps=0.51 for genes not converted. The average Pn=0.18 and Ps=0.44 for XI-MH63 gene conversion were significantly smaller than the average Pn=0.23 and Ps=0.49 for XI-MH63 genes with no conversion. XI-ZS97 gene conversion had average Pn=0.18 and Ps=0.45, which was significantly smaller than the average Pn=0.24 and Ps=0.49 for genes showing no conversion. We could not determine whether converted genes evolve slowly based on the paralogs themselves, since pairwise distances between paralogs are converted. However, Pn and Ps were slightly larger between orthologous gene pairs affected by gene conversion than between orthologs not showing gene conversion. This suggests that the orthologs in which gene conversion has occurred have evolved faster than those not affected by gene conversion.

**Table 3.** Nucleotide substitution rates of quartets in rice subspecies

| Paralog | XI-MH63 |  |  | XI-ZS97 |  |  | GJ |  |  |
| --- | --- | --- | --- | --- | --- | --- | --- | --- | --- |
|  | Pn | Ps | Pn/Ps | Pn | Ps | Pn/Ps | Pn | Ps | Pn/Ps |
| Gene conversion | 0.180 | 0.444 | 0.405 | 0.179 | 0.448 | 0.400 | 0.198 | 0.456 | 0.434 |
| No gene conversion | 0.234 | 0.486 | 0.481 | 0.235 | 0.486 | 0.484 | 0.253 | 0.509 | 0.497 |
| P-value | $1.82\times 10^{-18}$ | $8.73\times 10^{-9}$ | - | $7.81\times 10^{-20}$ | $1.25\times 10^{-7}$ | - | $3.71\times 10^{-26}$ | $8.00\times 10^{-20}$ | |
| Orthologs | XI-MH63 vs. GJ |  |  | XI-ZS97 vs. GJ |  |  | XI-MH63 vs. XI-ZS97 |  |  |
|  | Pn | Ps | Pn/Ps | Pn | Ps | Pn/Ps | Pn | Ps | Pn/Ps |
| Gene conversion | 0.049 | 0.076 | 0.645 | 0.053 | 0.085 | 0.624 | 0.055 | 0.100 | 0.550 |
| No gene conversion | 0.023 | 0.036 | 0.639 | 0.025 | 0.038 | 0.658 | 0.014 | 0.022 | 0.636 |
| P-value | $5.61\times 10^{-19}$ | $9.63\times 10^{-23}$ | - | $1.76\times 10^{-21}$ | $1.23\times 10^{-27}$ | | $6.25\times 10^{-20}$ | $2.58\times 10^{-31}$ | - |

We used Ps and Pn for determining whether gene conversion was affected by evolutionary selection pressure. The ratio of Pn/Ps reflects the selection pressure between gene pairs during evolution. We compared the Pn/Ps ratio between genes subjected to conversion and those with no conversion. The average Pn/Ps ratio for XI-MH63 gene conversion was 0.41, and the average Pn/Ps ratio of non-converted paralogs was 0.48. This indicates that converted genes were subject to purifying selection (**Table 3**). The Pn/Ps ratios for gene conversion in XI-ZS97 and GJ were also smaller than those for non-converted genes. The selection pressure for gene conversion or no gene conversion did not change much. However, there was not much difference in the selection pressure between orthologous gene pairs with and without gene substitution. No evidence suggests a change in selection pressure of converted genes.

### On-going gene conversion and function

Some duplicated genes are preferentially converted. We performed Gene Ontology (GO) analysis to relate duplicated genes to biological functions. The GO analysis revealed that some genes with specific functions may be preferred for conversion, while gene conversion of some functional genes is avoided (**Supplemental Figures 4-6**; **Supplemental Table 8**). We analyzed 761, 910, and 912 duplicated genes with gene conversion and 5,262, 5,224, and 5,135 duplicated genes without gene conversion in GJ, XI-MH63, and XI-ZS97, respectively. Genes involved in functions associated with large numbers of genes (catalytic activity, metabolic process) were biased toward gene conversion in the three genomes. By contrast, some genes associated with functions encoded by few genes (protein-containing complex, transporter activity) might have avoided gene conversion.

GO analysis of duplicated genes with and without gene conversion suggested that genes associated with functions encoded by a large number of genes are more biased towards gene conversion (**Table 4**). Four secondary-level terms were significantly enriched at the level of molecular function and biological processes, and accounted for about 30% of the corresponding gene sets. For example, the number of catalytic activity genes and metabolic process genes in the three genomes in which gene conversion occurred (31.4% - 37.7%) was significantly more than that in which no gene conversion occurred (26.6% - 30.6%) (P-value < 0.01). Similarly, binding genes and cellular process genes showed higher gene conversion (27.4% - 39.9%) than duplicated genes without gene conversion (24.6.6% - 38.4%), suggesting that they are more likely to be converted.

**Table 4.** Function comparison of genes subjected to conversion or not in GJ, XI-MH63, and XI-ZS97.

| GO level2 | GJ |  | XI-MH63 |  | XI-ZS97 |  |
| --- | --- | --- | --- | --- | --- | --- |
|  | cv vs. non-cv<sup>1</sup> | P-value | cv vs. non-cv<sup>1</sup> | P-value | cv vs. non-cv<sup>1</sup> | P-value |
| Catalytic activity | 31.4:25.5 | 0.001 | 32.5:26.4 | <0.001 | 33.6:26.1 | <0.001 |
| Binding | 38.8:35.5 | 0.083 | 39.9:37.7 | 0.199 | 39.6:38.1 | 0.412 |
| Metabolic process | 37.7:27.7 | <0.001 | 36.8:29.6 | <0.001 | 37.4:29.4 | <0.001 |
| Cellular process | 28.9:24.0 | 0.003 | 28.0:26.2 | 0.262 | 27.4:25.8 | 0.32 |

Note: <sup>1</sup>Proportion of converted genes vs. proportion of non-converted genes.

### Evolution and conversion of NBS-LRR genes

Rice diseases caused by various pathogens are one of the most serious constraints in global rice production (Divya et al., 2014). Disease resistance genes play a very important role in the evolution of plant genomes and are one of the indispensable families of genes for survival of plants under natural selection (Keen, 1992, Bertioli et al., 2016). We therefore identified 1,697 NBS-LRR (nucleotide binding site-leucine rich repeat) resistance genes in the three genomes (**Supplemental Table 9**). Among these, we identified 462 NBS-LRR genes in GJ, less than in XI-MH63 (644) and XI-ZS97 (591). The NBS-LRR genes were unevenly clustered on the chromosomes of the three genomes. The density on chromosome 11 was the highest, as confirmed in previous studies (Zhang et al., 2014, Stein et al., 2018). We found 113 (24.46%), 126 (21.32%), and 181 (28.11%) NBS-LRR genes on chromosome 11 of GJ, XI-MH63, and XI-ZS97, respectively. There were more NBS-LRR genes on chromosome 11 than on the other chromosomes (3.68% - 10.66%).

GO analysis of NBS-LRR genes in the genomes revealed enrichment mainly in terms associated with molecular function and biological process (**Supplemental Figure 7**). In GJ, XI-MH63, and XI-ZS97, 97%, 91.1%, and 93.1% of genes, respectively, were involved in binding (P-value=0.01) (**Supplemental Table 10**). Therefore, the NBS-LRR genes may be associated with the molecular function of binding and might be biased toward the occurrence of gene conversion. Polyploidization may also result in expansion of NBS-LRR genes, with ectopic recombination causing the NBS-LRR genes to further undergo a birth-to-death process. Evolutionary analysis of the NBS-LRR genes revealed 25, 67, and 39 young genes with *Ks* < 0.1 in the three genomes (**Figure 6A-C**). Most of the NBS-LRR genes were generated after the divergence of rice subspecies, and clusters of young NBS-LRR genes were found on chromosomes 2 and 11. These NBS-LRR genes showed a pattern of proximal localization and young origin in the three genomes, as well as similarity in gene conversion. We found a positive correlation between NBS-LRR genes and converted genes in regions with more than 1% of the NBS-LRR genes in the three genomes. This suggested that during their evolution, NBS-LRR genes might have had many chances to interact with one another, leading to gene conversion. (**Figure 6D**).

**Figure 6.**
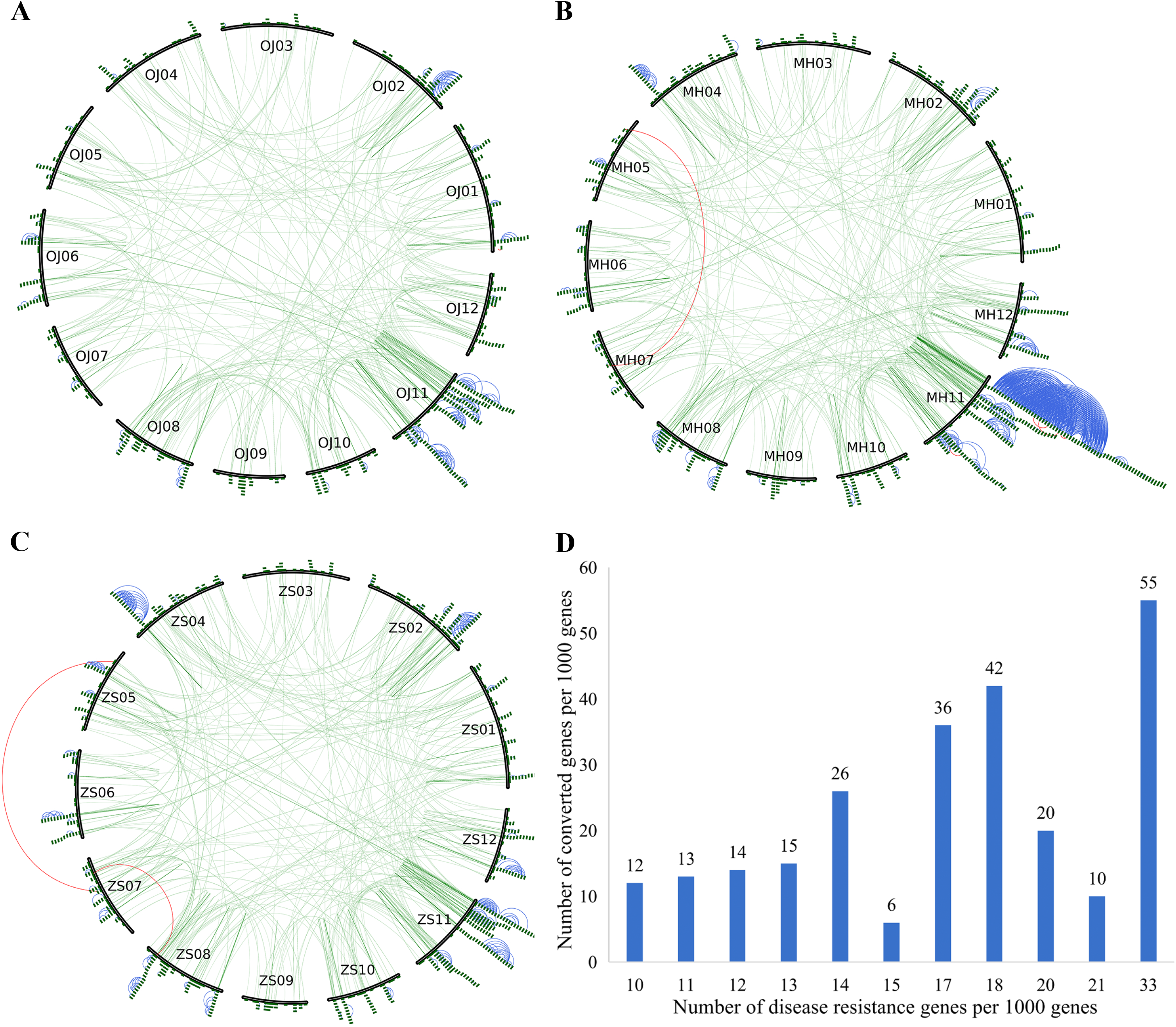
NBS-LRR gene amplification model in three rice subspecies genomes. (A-C) Distribution of NBS-LRR genes on 12 chromosomes in GJ, XI-MH63, and XI-ZS97. Green curved lines within the inner circle connect homologous pairs of NBS-LRR genes on the 12 chromosomes. Green blocks indicate NBS-LRR genes; red lines between NBS-LRR genes indicate *Ks* < 0.1, yellow lines indicate 0.1 < *Ks* < 0.2, and blue lines indicate *Ks* < 1. (D) Relationship between NBS-LRR genes and gene conversion in regions with more than 1% of the NBS-LRR genes in the three genomes.

## Discussion

### On-going conversion between duplicated genes

Recombination between neo-homologous chromosome pairs or homologous chromosome pairs resulting from WGD has existed throughout a long evolutionary history, generated a large number of chromosomal rearrangements (Murat et al., 2010, Bowers et al., 2003, Murat et al., 2014). This recombination can persist for a long time, maybe even hundreds of millions of years (Wicker et al., 2015). Previous studies have illustrated that many duplicated genes from WGD events about 100 mya are affected by illegitimate recombination and gene conversion (Jacquemin et al., 2009, Jacquemin et al., 2011). In some genomic regions, this effect persists for millions of years, especially on chromosomes 11 and 12 of rice (Wang et al., 2007). We used new, high-quality genomic data to analyze the genome sequences of GJ, XI-ZS97, and XI-MH63, revealing the level and pattern of gene conversion in all homologous genes of modern crop rice during domestication and improvement. Study of gene conversion after the divergence of rice subspecies and after the divergence of the two XI varieties revealed a shared region of gene substitution between XI-MH63 and XI-ZS97. This suggests that gene conversion may be on-going for a long time in the evolution of species and continue to provide a driving force in genome evolution and genetic innovation.

### Gene conversion has contributed to cultivated rice divergence

Gene conversion is the result of recombination. Classical theoretical studies point out that recombination accelerates mutation (Koszul and Fischer, 2009, Jacquemin et al., 2011). Gene conversion may therefore play an important role in recombination, and we used the results of the new data analysis to further confirm this conclusion. We identified that the *Ks* between orthologous genes showing gene conversion was significantly smaller than that of orthologous genes without conversion. This suggests that genes having undergone gene conversion may have evolved more rapidly, which has been demonstrated by previous studies (Chen et al., 2007, Wang and Paterson, 2011). Gene conversion is one of the major mutational mechanisms in the evolution of species. Gene conservation can provide opportunities for gene conversion (Cossu et al., 2017). Our results showed that 46% of ancient gene conversions may have again undergone gene conversion more recently after the divergence of rice subspecies. Gene conversion is an accelerating force in the genetic evolution of mutations. After gene conversion, these genes restart the evolutionary process and accelerate the divergence of rice subspecies.

### Gene conversion and chromosome rearrangement

Our results showed that the degree of chromosome rearrangement and gene conversion rate are positively correlated. However, gene conversion is not necessary for the survival of the species, as most grass species have undergone massive chromosome rearrangements (Murat et al., 2010, Wang et al., 2019). Previous reports suggest that the occurrence of a large inversion in the short arm before the rice–sorghum divergence may suppress gene conversion, with the lowest rate of gene conversion occurring between chromosomes 1 and 5 in rice (Wang et al., 2009, Paterson et al., 2009). However, we did not find the lowest rates of gene conversion in the three genomes of rice subspecies between chromosomes 1 and 5, possibly because chromosome recombination may be stage-specific. Shorter homoeologous regions are a modern state resulting from historical evolution. We found more chromosomal rearrangements in XI than GJ, which may lead to gene loss, and relatively more gene conversion. Chromosome rearrangement might result in gene loss and thus provide conditions for on-going gene conversion. Chromosome rearrangement might have directly contributed to restriction of recombination/conversion between homoeologous regions.

### Why is a donor usually a donor?

Gene conversion is to copy one gene sequence from a donor locus to a receptor locus (Harpak et al., 2017). Analyzing the scale of gene conversion helps to illuminate the mechanism of gene conversion (Cossu et al., 2017). We found that independent conversions that have survived (so far) in different lineages have often used the same genes as donors. It seems improbable to attribute this to selection, noting that the donor and acceptor copies have coexisted in the genome for 100 million years. A more plausible explanation is that one gene copy has some ‘privileged’ nature over the other. This could be genetic or epigenetic. If one copy or its neighboring region possesses mutations or epigenetic changes, the other copy might be more likely to act as a donor, helping to reinstate intactness. Moreover, some homologous chromosomal segments also seem to be preferential donors rather than acceptors. Mechanisms underlying these biases remain unknown, but an exciting future investigation will be to explore epigenetic phenomena such as have been suggested to influence patterns of gene retention/loss along chromosome segments (Woodhouse et al., 2010).

### Gene conversion and function

Gene conversion leads to genes similar or even identical in sequence. The analysis above indicates that large gene families may be more susceptible to gene conversion. Duplicated copies may neutralize the presence of putative mutations, providing an opportunity for functional innovation (Daugherty and Zanders, 2019). Rather than being a conservative factor among different genotypes, gene conversion accelerates divergence (Wang et al., 2011). Gene conversion has been used to explain the evolution of large gene families, such as NBS-LRR genes and rRNA genes, which typically have dozens of copies on chromosomes (Okuyama et al., 2011, Nawrocki and Eddy, 2013, Rooney, 2004). Extensive analysis has shown that the evolution of functional genes that are members of large families may often be accompanied by strong purifying selection. Until 1990, most multigene families were thought to have coevolved with related homologous genes through gene conversion (Godiard et al., 1994).

Evolution of the NBS-LRR gene family, rRNA gene family, and some other highly conserved gene families may be consistent with this conclusion. For these families, most genes are usually extremely similar. However, the evolution of other gene families may be better explained by the birth-and-death model. New genes are created through gene duplication, and some genes remain in the genome for a long time while others may be lost (Finet et al., 2019).

## Materials and Methods

### Sequence data

Genomic sequence data for XI-MH63 and XI-ZS97 were obtained from the GenBank database (https://www.ncbi.nlm.nih.gov/). Genomic data for GJ ‘Nipponbare’ and *Arabidopsis thaliana* were downloaded from genome databases Gramene (http://www.gramene.org/) and TAIR (https://www.arabidopsis.org/), respectively.

### Detection of duplicated segments and homologous gene quartets

BLASTP (Camacho et al., 2009) was used to search for intragenomic and intergenomic homology of protein sequences (E < 1e-5). ColinearScan (Wang et al., 2005) was used to analyze colinear regions based on gene homology predictions, and the significance of colinearity was tested. Colinear intragenomic and intergenomic chromosome fragments were inferred from analysis of homologous genome structures, and homologous and colinear genes were determined. Blocks of homologous genome structure within and between rice subspecies were also deduced. These blocks might represent paralogs produced by WGD events in the common ancestor or orthologs caused by species divergence. To determine homology and colinearity between chromosomes, genes in large gene families were removed from the ColinearScan analysis. Therefore, to obtain more complete homology information within genomes, further bidirectional best BLASTP homology searches were performed on the three genomes. Gene quartets were inferred from intragenomic and intergenomic paralogs and orthologs.

### Inference of gene conversion

To infer possible gene conversion between paralogs, ClustalW (Larkin et al., 2007) comparison of the quartets identified between any two genomes was performed. Highly divergent sequences were removed to eliminate potential problems created by inferring gene conversion from unreliable sequences. Quartets showing gaps in the pairwise alignments exceeding 50% of the alignment length, or with amino acid identity between homologous sequences of less than 40% were removed.

#### *Whole-genome conversion* (WCV) inference

Since paralogous genes arise before species divergence, the similarity between orthologous gene pairs in two species should be higher than the similarity between paralogous gene pairs. However, gene conversion events change the similarity between gene pairs. The first whole-genome conversion inference method (WCV-I) used was based on studying the homology relationship between genomes, using *Ks* value as a similarity measure. The *Ks* values between paralogous and orthologous gene pairs were used to infer possible gene conversion, and 1000 bootstrap tests were performed on all gene trees in which gene conversion occurred to obtain the confidence level for each gene (Wang et al., 2009, Wang and Paterson, 2011). The second whole-genome conversion inference method (WCV-II) calculated the ratio of amino acid locus identity between homologous gene pairs, and compared point-by-point homology between paralogous gene pairs and between orthologous gene pairs. These sequences were used to infer possible changes to evolutionary tree topology, depending on whether the paralogous genes were more similar to each other than orthologous genes (Wang et al., 2009). This is a strict criterion, as paralogs were produced at least 100 mya from a WGD, whereas orthologs have diverged more recently. Instead of using *Ks* values as a metric here, identical sites between homoeologous sequences were calculated directly. The similarity between sequences representing different rice subspecies is often very high, as in a previous study of hexaploid wheat (Liu et al., 2020).

#### *Partial-gene conversion* (PCV) inference

Quartets were used to identify possible gene conversion among partial gene sequences that may occur after species divergence. A combination of dynamic planning and phylogenetic analysis was used to document the differences between two aligned bases from paralogous and orthologous genes for each genome. In averaged distance arrays, the paralogs in each species should be more distant if no PCV was involved. Bootstrap frequency was obtained by repeating the 1000 bootstrap tests to identify high-scoring segments with shorter lengths and smaller scores. After masking some of the larger fragments, a recursive procedure revealed shorter high-scoring fragments, which helped to reveal genes affected by multiple gene conversion events (Wang et al., 2009).

### GO enrichment analysis

The GO data search software InterProScan (Jones et al., 2014) was used to determine whole-genome GO functional annotation. GO annotation results of the gene sets were compared and plotted using the online visualization tool WEGO (Ye et al., 2018) to visualize the distribution of functional genes and trends. The significance of the enrichment of GO-annotated genes was explained using calculated P-value.

### Identification of disease-resistance genes

The comparison software HMMscan (Eddy, 2011) was used to identify NBS-LRR domains in the whole genomes of GJ, XI-MH63, and XI-ZS97, and NBS-LRR gene set A was obtained. The whole genome of the model organism *Arabidopsis thaliana* was searched for the NB-ARC domain (PF00931) using HMMsearch (Eddy, 2011) to identify NBS-LRR domains with E-value of 1e-10. After obtaining the NBS-LRR genes of *Arabidopsis thaliana*, BLASTP was used to compare these sequences with the whole genomes of GJ, XI-MH63, and XI-ZS97. Genes with a score value of > 150 and E-value > 1e-10 were designated NBS-LRR gene set B of the rice subspecies. Genes present in both gene sets A and B were identified as NBS-LRR genes in the three genomes.

### Accession Numbers

Sequence data from this article can be found in Materials.

## Supplemental Data

**Supplemental Text** Gene conversion and occurrence patterns.

**Supplemental Figure 1.** Distribution of amino acid identity between duplicated genes in *Oryza* subspecies genomes.

**Supplemental Figure 2.** Distribution of synonymous nucleotide substitution percentage (Ps) between syntenic paralogs in duplicated blocks of *Oryza* subspecies genomes.

**Supplemental Figure 3.** Relationship between length of blocks on each chromosome and rate of gene conversion.

**Supplemental Figure 4.** Histogram of Gene Ontology (GO) statistics for duplicated genes with and without gene conversion in GJ.

**Supplemental Figure 5.** Histogram of Gene Ontology (GO) statistics for duplicated genes with and without gene conversion in XI-MH63.

**Supplemental Figure 6.** Histogram of Gene Ontology (GO) statistics for duplicated genes with and without gene conversion in XI-ZS97.

**Supplemental Figure 7.** Histogram of Gene Ontology (GO) statistics of NBS-LRR genes in GJ, XI-MH63 and XI-ZS97.

**Supplemental Table 1.** Number of homologous genes and blocks in GJ, XI-MH63, and XI-ZS97.

**Supplemental Table 2.** Identified quartets and gene conversion in GJ, XI-MH63, and XI-ZS97.

**Supplemental Table 3.** Gene conversion of quartets in the three rice subspcies genomes.

**Supplemental Table 4.** Homology of donor locus and acceptor locus in gene conversion.

**Supplemental Table 5.** Distribution of paralogs and gene conversion GJ, XI-MH63, and XI-ZS97.

**Supplemental Table 6.** Relationship between the block number and the gene conversion rate in GJ, XI-MH63, and XI-ZS97.

**Supplemental Table 7.** Relationship between the block length and the gene conversion rate in the three rice subspecies genomes.

**Supplemental Table 8.** GO analysis of gene conversion and non-gene conversion in GJ, XI-MH63, and XI-ZS97.

**Supplemental Table 9.** NBS-LRR gene counts by chromosome in GJ, XI-MH63, and XI-ZS97.

**Supplemental Table 10.** GO annotation analysis of NBS-LRR genes in GJ, XI-MH63, and XI-ZS97.

## Competing interests

The authors declare no competing financial interests.

## Acknowledgments

We appreciate the financial support from the China National Science Foundation (3151333 to JW), Natural Science Foundation of Hebei Province (C20209064 and C2015209069 to JW).

